# Data Representation in the DARPA SD2 Program

**DOI:** 10.1101/2021.09.17.460644

**Authors:** Nicholas Roehner, Jacob Beal, Bryan Bartley, Richard Markeloff, Tom Mitchell, Tramy Nguyen, Daniel Sumorok, Nicholas Walczak, Chris Myers, Zach Zundel, James Scholz, Benjamin Hatch, Mark Weston, John Colonna-Romano

## Abstract

Modern scientific enterprises are often highly complex and multidisciplinary, particularly in areas like synthetic biology where the subject at hand is itself inherently complex and multidisciplinary. Collaboration across many organizations is necessary to efficiently tackle such problems [6, 15], but remains difficult. The challenge is further amplified by automation that increases the pace at which new information can be produced, and particularly so for matters of fundamental research, where concepts and definitions are inherently fluid and may rapidly change as an investigation evolves [7].

The DARPA program Synergistic Discovery and Design (SD2) aimed to address these challenges by organizing the development of data-driven methods to accelerate discovery and improve design robustness, with one of the key domains under study being synthetic biology. The program was specifically organized such that teams provided complementary types of expertise and resources, and without any team being in a dominant organizational position, such that subject-matter investigations would necessarily require peer-level collaboration across multiple team boundaries. With more than 100 researchers across more than 20 organizations, several of which ran experimental facilities with high-throughput automation, participants were forced to confront challenges around effective data sharing.

The default architecture for scientific collaboration is essentially one of anarchy, with ad-hoc bilateral relations between pairs of collaborators or experimental phases (Figure 1(a)). This was by necessity the case during early phases of the SD2 program as well, in which incorporating new tools into pipelines was ad-hoc and time-consuming, and data was generally disconnected from genetic designs and experimental plans. The other typical approach for collaboration is one of “command and control”, in which a dominant organization determines the data sharing content and format for all participants (Figure 1(b)). This can be efficient, but tends to be limited in flexibility and extensibility, rendering it unsuitable for research collaboration, as indeed was found when we attempted this approach during the first year of the SD2 program. We addressed these problems with the application of distributed standards to create a “flexible rendezvous” model of collaboration (Figure 1(c)), enabling information flow to track evolving collaborative relationships, improving the sharing and utility of information across the community and supporting accelerated rates of experimentation.

## 2 APPROACH

The driving design philosophy behind our approach to user interaction in SD2 was to adapt representational tooling as closely as possible to existing tools and familiar interfaces, such as spreadsheets and word processing documents. Taking this approach allowed us to use and improve formal machine-readable representations for system integration while minimizing the amount that participating researchers needed to learn about the formal representations. The central set of standards thus formed a point of rendezvous between the various stakeholders interacting in different experimental roles, while still allowing each of these participants to continue working in their native idiom.

**Figure 1:**
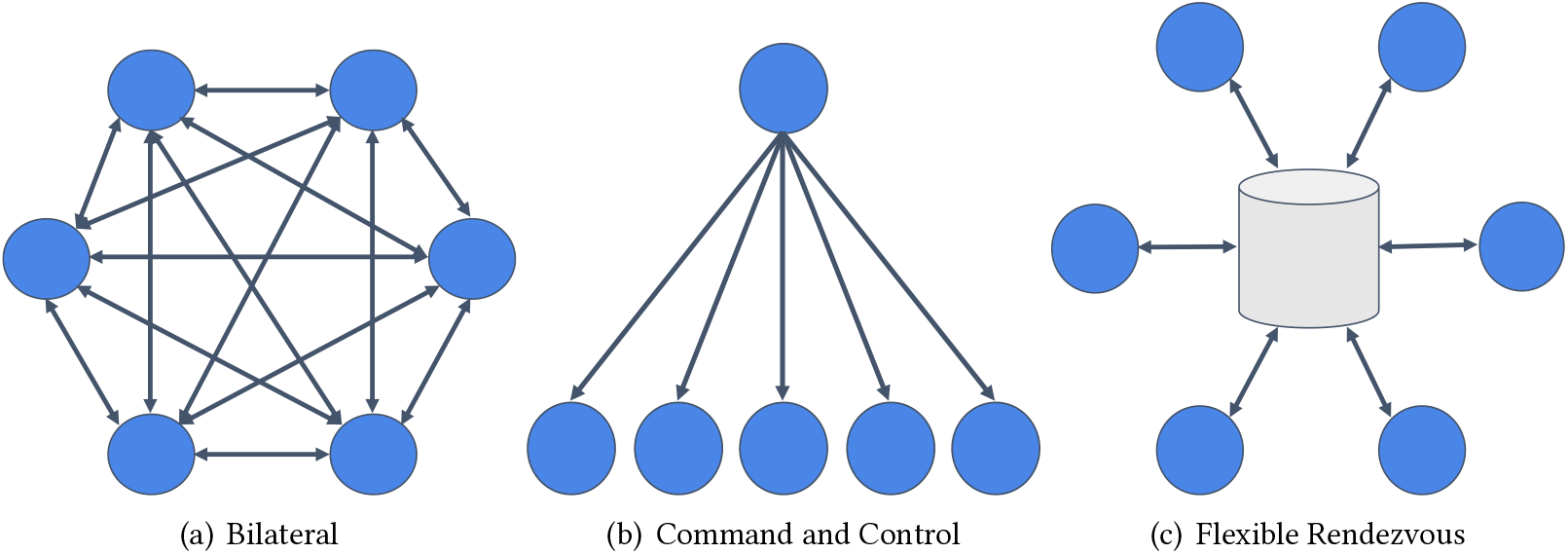
Architectures for data sharing: bilateral relations (a), command and control (b), and flexible rendezvous (c).

Specifically, the data sharing working group collaborated in the creation of several key advancements in standards and tooling that combine to create a comprehensive ecosystem of lightweight curation tools. These are:

- Advances in biological data representation in the form of enhancements to SBOL2 for comprehensive representation of the design-build-test-learn cycle [5, 8] and ultimately the development of SBOL3 [11], which in turn enabled us to accelerate the rate of data standard development and integration.
- A plugin interface to SynBioHub [12], which enabled rapid development of new functionality for data visualization, submission, and exchange [10].
- The SBOL Project Dictionary tool [4], which provides a Google Sheets interface for collective “just-in-time” harmonization of terminology across organizations, thus enabling metadata translation and data fusion. The Experimental Intent Parser tool [13], an extension of Google Docs that enabled biologists to design and launch automated experiments with an easy-to-use interface.
- The Open Protocol Interface Language (OPIL) [1], which enabled the laboratories executing experiments to share information about their protocols with experiment planning tools, thus better informing the investigators proposing experiments to execute.
- The SYNBICT tool [14], which enabled automated generation of improved annotations and extraction of functional models of biological designs from their sequences.
- The REDOER tool [2], which attempts to infer experimental design from collections of samples, enabling quality-control on automated experimentation.
- The Excel2SBOL conversion tool [9], which enabled an efficient workflow for producing build requests for genetic designs.

Collectively deployed in the architecture shown in Figure 2, these tools enabled a shift in the organization of experimental and informational workflows toward faster and more flexible execution, most notably in SD2 working groups that were working on challenge problems focused on the performance of genetic circuits in yeast, moving existing designs into novel chassis, and cell-free riboswitch design.

**Figure 2:**
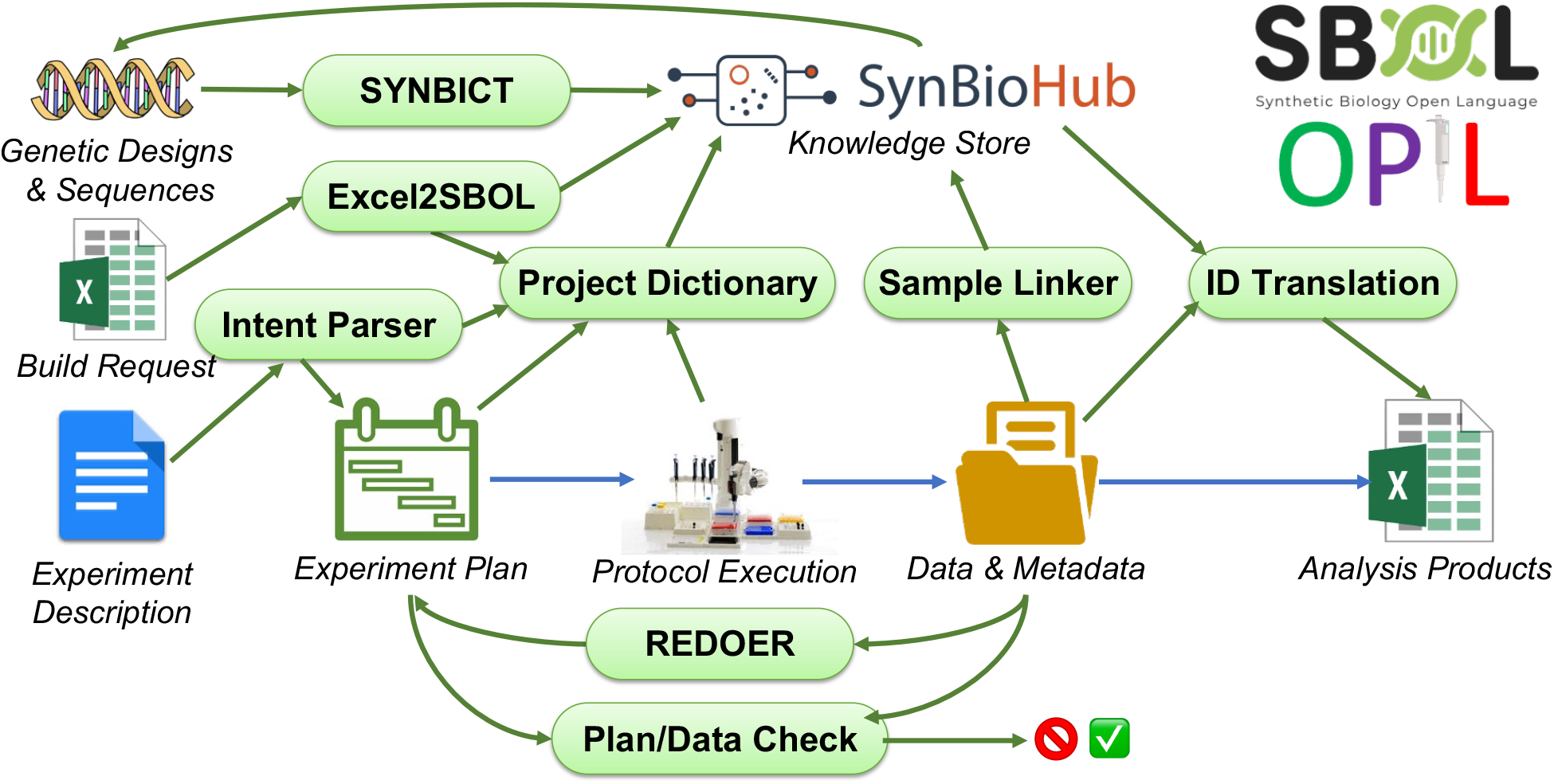
High-level diagram showing how data representation tools were deployed in the DARPA SD2 program with respect to inputs of designs, build requests, and experiments requests, and outputs of data, metadata, and analysis products. This diagram focuses specifically on the key representations (SBOL and OPIL) and representation-centric tooling, and not other aspects of supporting automation used in SD2.

Following this shift, program knowledge sharing expanded greatly. One key measure of knowledge sharing is the number of terms stored in the SBOL Project Dictionary, as each such term indicates a strain, reagent, genetic construct, parameter, or other similar item that is being communicated between collaborating organizations. We find that the number of terms stored in the SBOL Project Dictionary, expanded in close correlation with the increase in experiment tempo: Figure 3 shows the correlation between knowledge sharing and data production, with both moving much more quickly in the second half of SD2 after these tools began to be released.

Moreover, measurement of key knowledge collections shows that progress on tools correlates with increases in knowledge sharing and productivity. Figure 4 shows that knowledge expansions in SynBioHub are correlated with the dates of tool releases. In particular, key tool releases occurred around July 2018 (Project Dictionary), April 2019 (SYNBICT, REDOER, and Intent Parser), October 2019 (Experiment launches via Intent Parser), January 2020 (Excel2SBOL), April 2020 (SYNBICT libraries), and October 2020 (OPIL), and these are correlated with expansions in the number of SBOL Module relations, which are used in representing knowledge about circuits, and the number of `ModuleDefinition` relations, which are used in representing strains and reagents.

**Figure 3:**
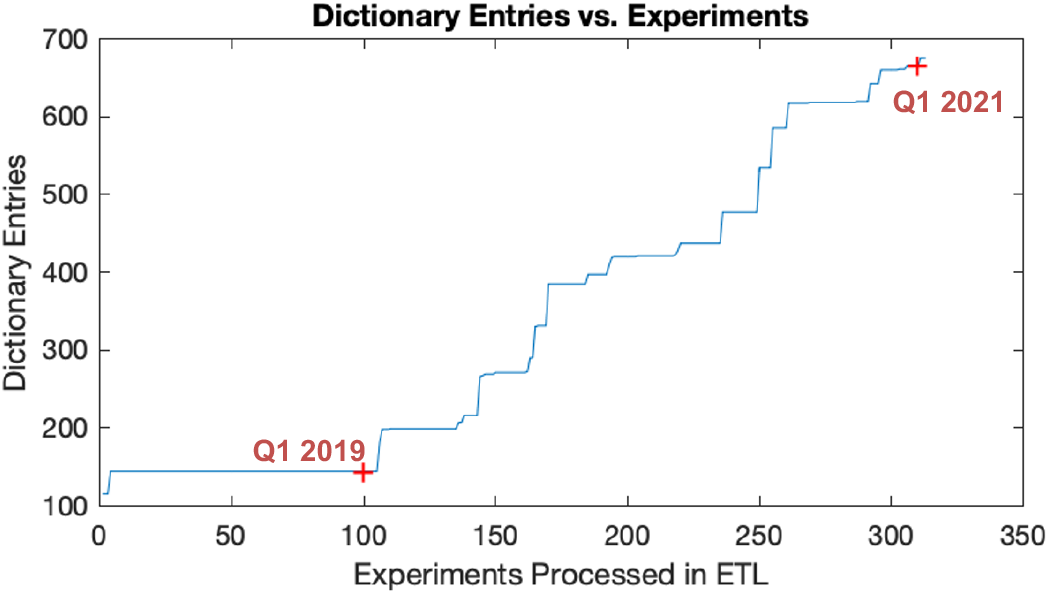
Increased knowledge sharing is correlated with overall rates of experimentation in SD2. Accumulation of shared knowledge, as measured by increased numbers of entries in the SBOL Project Dictionary, increased much more rapidly in the second half of the SD2 program, as did the rate at which experiments were run.

**Figure 4:**
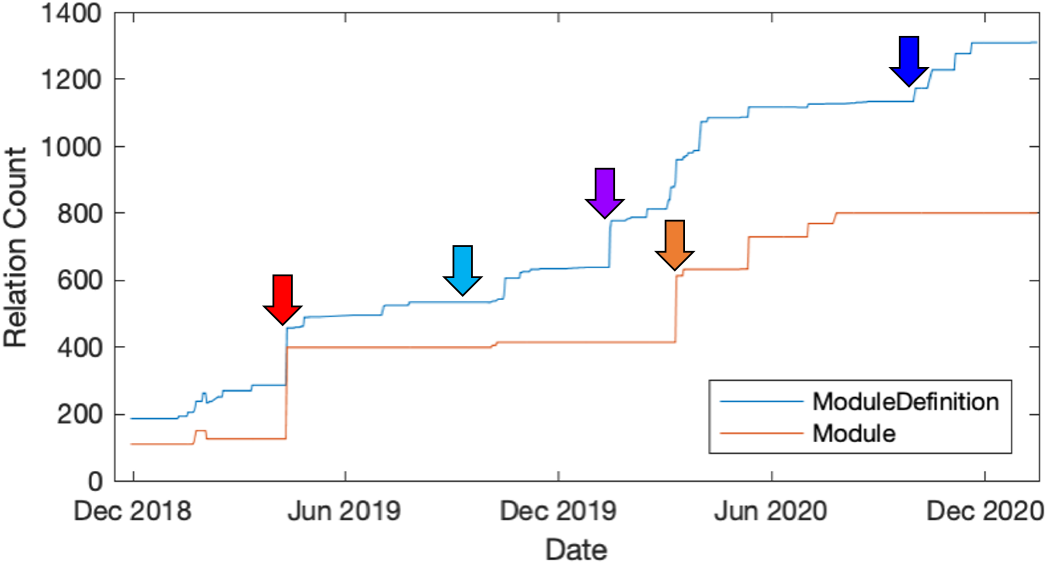
Introduction of specific tools is SD2 is correlated with expansions in key forms of shared knowledge. Arrows mark the approximate time of introduction of key advances in data representation tools: SYNBICT and Experimental Intent Parser (red), experiment launches via Experimental Intent Parser (light blue), Excel-to-SBOL (purple), SYNBICT libraries (orange), and OPIL (dark blue).

Finally, beyond these specific knowledge classes, the overall volume of knowledge systematized by the program is quite large as well. By the end of the period reported, the SD2 SynBioHub instance had a knowledge store of 22,872,306 triples, including 3,549,237 components, 15,884 modules, and 524,377 collections.

## 3 DISCUSSION

The transition partners that we have engaged with these tools see potential value in using the Synthetic Biology Open Language (SBOL) and associated tools with well-defined APIs as a standard that is relevant to many groups and has greater potential for sharing, greater levels of support, and more longevity. Partners can benefit from work by the supporting community and worry less about their data-sharing infrastructure losing “product support.” Transition partners also see value in standards and tools being free and open, which allows them to know what their data-sharing methods are doing, modify them if needed, interact with them easily via their own code, and not be limited by high commercial costs or number of licensed “seats.” With solutions like those that we developed in the SD2 program, they will be better able to keep track of information over time as new people come and go from labs, so that they can continue to build new knowledge on top of existing knowledge. They will also be better able to share design information and associated data with other research groups in a more consistent way, and be better able to take advantage of other groups’ designs and data, making their own engineering processes much faster.

Building on the success of data representation in the SD2 program, we recommend that future programs should support the development of standards and corresponding software infrastructure and also should support curators and the development of data repositories and curation tools. Similarly, just as many government funding agencies now require open access publications, government funding agencies should require funded science activities to use standard-enabled workflows and software to enhance data management and sharing, and should ensure that funding is specifically allocated for such activities. Looking farther ahead, we also see an opportunity for increased machine-readability of shared information to become a foundation for higher-level autonomy in scientific investigation [3], as well as for enhancing reproducibility through machine validation of scientific experiments and automation-assisted publication of experiments.

## 4 ACKNOWLEDGEMENTS

This work was supported by Air Force Research Laboratory (AFRL) and DARPA contracts FA8750-17-C-0184, FA8750-17-C-0229, FA8750-17-C-0231, and FA8750-17-C-0294. This document does not contain technology or technical data controlled under either U.S. International Traffic in Arms Regulation or U.S. Export Administration Regulations. Views, opinions, and/or findings expressed are those of the author(s) and should not be interpreted as representing the official views or policies of the Department of Defense or the U.S. Government.

